# An improved deep learning model for immunogenic B epitope prediction

**DOI:** 10.1101/2025.07.29.667398

**Authors:** Rakshanda Sajeed, Swatantra Pradhan, Rajgopal Srinivasan, Sadhna Rana

## Abstract

The recognition of B epitopes by B cells of the immune system initiates an immune response that leads to the production of antibodies to combat bacterial and viral infections. Computational methods for predicting the epitopes on antigens have shown promising results in the development of subunit vaccines and therapeutics. Recently, the use of protein language models (pLMs) for epitope prediction has led to a substantial increase in prediction accuracy. However, further improvements in precision are necessary for practical applications. Here, we develop and evaluate a series of models using different combinations of features and feature fusion techniques on a curated independent test set. Our results show that the models that use protein embeddings along with structural features are better at predicting both linear and conformational B epitopes when compared to a baseline model that uses only protein embeddings as features. Additionally, we show that the embeddings of ESM-2, an evolutionary scale model, likely capture T-B reciprocity.

## 1 Introduction

B cell epitopes are special regions of an antigen sequence that are recognized by B cells and B cell receptors (BCR). The binding of an antigen epitope with B cell/BCR triggers a highly specific immune response that leads to the neutralization of pathogens like viruses, bacteria, or other foreign substances (Bahai et al., 2021). Thus, B epitope identification is crucial in the field of immunotherapy and vaccine design.

B-cell epitopes can be identified by different experimental techniques such as solving the 3D structure of antigen-antibody complexes, nuclear magnetic resonance spectroscopy, peptide library screening of antibody binding or performing functional assays in which the antigen is mutated and the interacting antibody-antigen is evaluated (Sanchez-Trincado et al., 2017). However, these methods are expensive, time-consuming and some require a high level of lab expertise. This is why the development of *in silico* tools has attracted a lot of attention. Recently, the field of computational epitope prediction has seen a surge in the use of advanced deep learning methods for improving the accuracy of B epitope predictions. BepiPred3.0, a sequence-based epitope prediction tool has leveraged evolutionary scale modeling (ESM-2) (Lin et al., 2022) to improve the prediction accuracy for both linear and conformational epitope predictions. DiscoTope-3.0 (Høie et al., 2024) maintains high predictive performance across antigen structures solved experimentally as well as those predicted using AlphaFold2.0 (Jumper et al., 2021), thus alleviating the need for experimental structures. Similarly, SEMA 2.0 (Ivanisenko et al., 2024) has shown improved prediction accuracy using ESM and SaProt, a structure-aware protein language model (Su et al., 2024). These current generation tools using language models and deep learning models such as AlphaFold2.0 have shown significant improvement in prediction as compared to other conformational epitope prediction tools from previous generation such as, BepiPred2.0 (Jespersen et al., 2017), Ellipro (Ponomarenko et al., 2008), Discotope-2.0 (Kringelum et al., 2012), SEPPA3.0 (Zhou et al., 2019), Epitope-3D (da Silva et al., 2022), which use extensive feature engineering and include structural, sequential and evolutionary features of the protein for training the models.

In this study, we aim to enhance B epitope prediction by integrating structural features with those derived from protein language models. We evaluated different approaches for feature combination, including naive concatenation as well as deep transformation and concatenation methods, to effectively merge protein embeddings with structural characteristics within our models. Our findings indicate that, on a curated independent test set, combining structural features with protein embeddings using deep transformation and concatenation technique delivers superior results compared to naive feature concatenation or a baseline model that relies solely on protein embeddings. Our best model with combined features outperforms other state-of-the-art conformational epitope predictors on most of the metrics. We also show that structural features boost the prediction of linear epitopes as well. Previous studies have included T-B reciprocity (Zhu et al., 2022) as a feature for improved epitope prediction. Attention analysis of antigen sequences indicated that ESM-2 embeddings likely capture T-B reciprocity, as a substantial proportion of high-scoring B epitopes are highly attended by T epitopes.

## 2. Materials and Methods

### 2.1 Training and test data for conformational epitopes

The IEDB-3D (Ponomarenko et al., 2011) full-bcr assay dataset was the basis for our training and evaluation process (available at http://www.iedb.org/database_export_v3.php, date of download, 5th August 2024). The dataset consisted of 4554 entries with antigen information, antibody information, host information and the experimental information. To ensure a high-quality data for training, we retained only antigen-antibody crystal structures with resolution better than 3Å and the entries with protein epitopes, reducing the number of data points from 4554 to 1827. We also removed antigen sequences shorter than 60 amino acids, resulting in 1491 unique antigen-antibody complexes from PDB.

To reduce bias in the data due to similar sequences, we performed a redundancy check on antigen sequences using a 50% identity threshold (Clifford et al., 2022) with the MMSeqs2 (Steinegger and Söding, 2017) clustering tool. For each cluster, only the protein sequence with the highest number of residues in the epitope region was retained, resulting in 901 antigen-antibody crystal structures. The antigen sequences from these complexes were provided as input to ESMFold (Lin et al., 2022) to generate apo, or unbound, structures of the antigens. This approach aimed to limit potential data leakage by avoiding the influence of antibody presence during epitope prediction, facilitating the identification of putative epitope regions without antibody information. In the final antigen sequence data, positive datapoints refer to residues experimentally characterized as epitopes, while the remainder were considered non-epitopes or negative datapoints. These 901 complexes were partitioned into 5 groups with 180 complexes in each. In each group we balanced the positive and negative datapoints.

We evaluated our models on a non-redundant independent test data created from SabDAb, the Dset_anti antigen dataset (Hou et al., 2021) consisting of 280 antigen-antibody complexes. To compare with known epitope prediction tools BepiPred2.0 (Jespersen et al., 2017), BepiPred3.0 (Clifford et al., 2022), DiscoTope2.0 (Kringelum et al., 2012), DiscoTope3.0 (Høie et al., 2024) and SEMA2.0 (Ivanisenko et al., 2024) we checked the similarity of their training sets with our test set at 25% sequence identity thresholds. After executing redundancy checks (25% identity cut off) to prevent data leakage with our training set and the training sets of other predictors that were used for model comparison, we were left with 45 antigen-antibody complexes in our final independent evaluation dataset.

### 2.2 Training and test data for linear epitopes

We extracted the linear B-cell epitope data from IEDB. We selected antigens that were annotated with the UniProt ID so that they could be mapped to a PDB (apo) structure. There were 1184 datapoints with the parent protein information. We also followed the other filtration steps, such as considering the resolution of the structure (better than 3Å) and a redundancy check (25% identity threshold) for the antigens. Finally, we were left with 438 antigens, which were partitioned into train, validation and test sets consisting of 339, 27 and 72 antigens respectively.

### 2.3 Structural Feature Generation

We computed key structural features for the protein sequences. These features are contact number (CN) (Yuan, 2005), protrusion index (PI) (Xia et al., 2010) and half sphere exposure (HSE) (Sweredoski and Baldi, 2008). The contact number was computed as the number of residues whose Cβ atoms were present within a distance threshold (10Å) of the residue of interest. Protrusion index is a geometric structural feature which estimates the extent to which the residue is protruding from the surface of the protein complex. The number of heavy atoms (N_atoms_) within a fixed distance R (10Å) was calculated. Then, N_atoms_ is multiplied by the mean atomic volume which is the volume occupied by the protein within the sphere, V_int_ which is 20.1 ± 0.9Å^3^. The difference between the volume of the sphere and V_int_ is given by V_ext_. The protrusion index is then calculated as V_ext_/V_int_. Half Sphere Exposure measures the solvent exposure of a residue in the protein and is calculated by counting the amino acid neighbours within the two half spheres of a certain radius (usually 12Å) around the amino acid residue. This shows how buried the residue is in the protein. For this, the BioPDB (“Bio.PDB package — Biopython 1.75 documentation,” n.d.) (Cock et al., 2009) module HSExposure was used. The HSE is calculated for the residues of the known protein chain around a radius of our requirement.

### 2.4 Model Architecture

Protein embeddings and structural features represent an epitope from different perspectives. To improve epitope prediction, we used two feature fusion techniques: naive vector concatenation, and deep transformation and concatenation to obtain an informative representation of an epitope. We developed two neural network models for B-cell epitope classification, both using the ESM-2 650M model as a feature extractor to generate a 1280-dimensional vector for each datapoint. The two architectures are as follows:1. Architecture I or Naive Vector Concatenation: In this architecture (Figure 1), we performed a simple concatenation of the ESM-2 embeddings with the different combinations of structural features to obtain a single representation of the input. This way of combining different feature sets is commonly seen to work well in all settings. This kind of approach ignores the fact that embeddings and structural features are from different spaces. The final representation of the input with 1280(+1/2/3) (based on the number of structural features considered) features was given as input to a 4-layer neural network where the weights of the neural network were learned to do the classification of the residue into one of the two classes epitope vs non-epitope.

**Figure 1.**
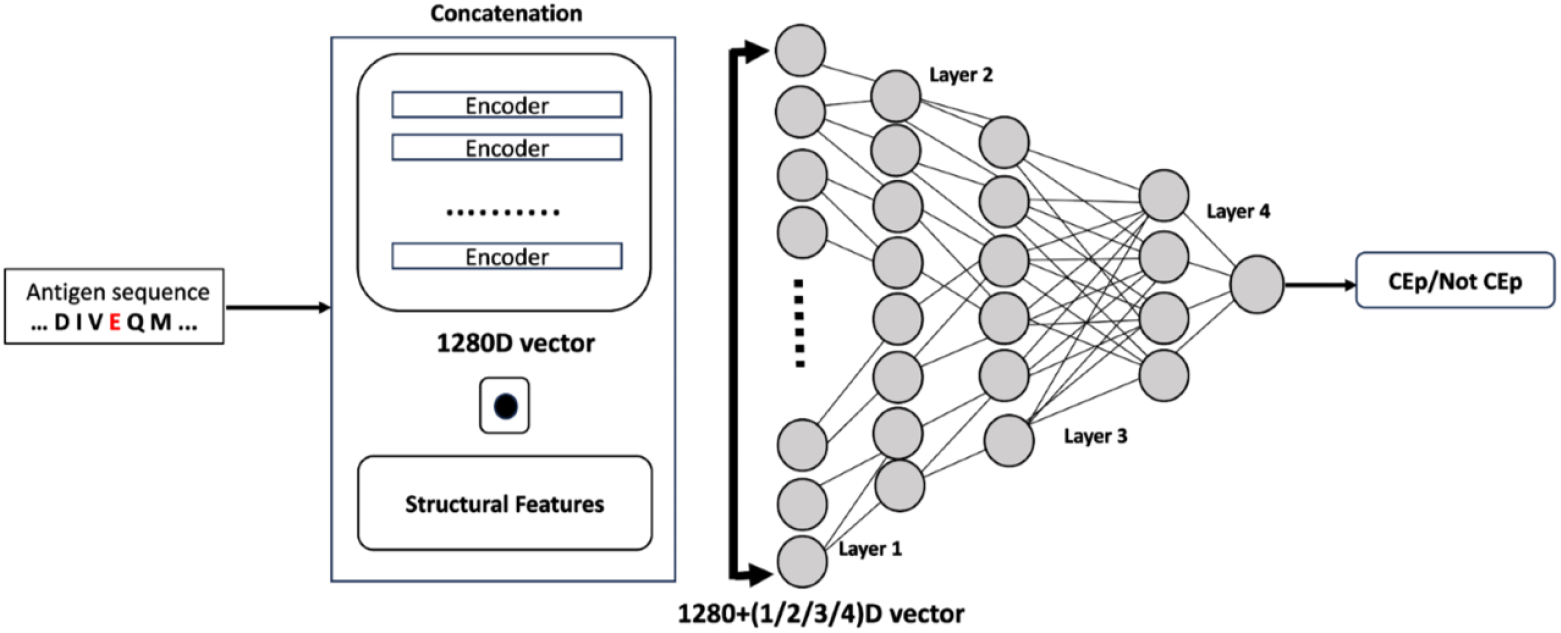
Model architecture I shows naive concatenation of structural features and ESM-2 embeddings at the input layer, processes them through four hidden layers, and outputs a probability score to classify residues as epitopes or non-epitopes.

2. Architecture II or deep transformation and concatenation of multi modal Features: In this architecture (Figure 2), we learn different weights as well as common representations for protein embeddings and structural features before concatenation. To achieve this, the ESM-2 embeddings were transformed through the first 4 layers of the neural network, then the transformed feature vector was concatenated with different combinations of structural features which was then given as input to the final layer of the NN.

**Figure 2.**
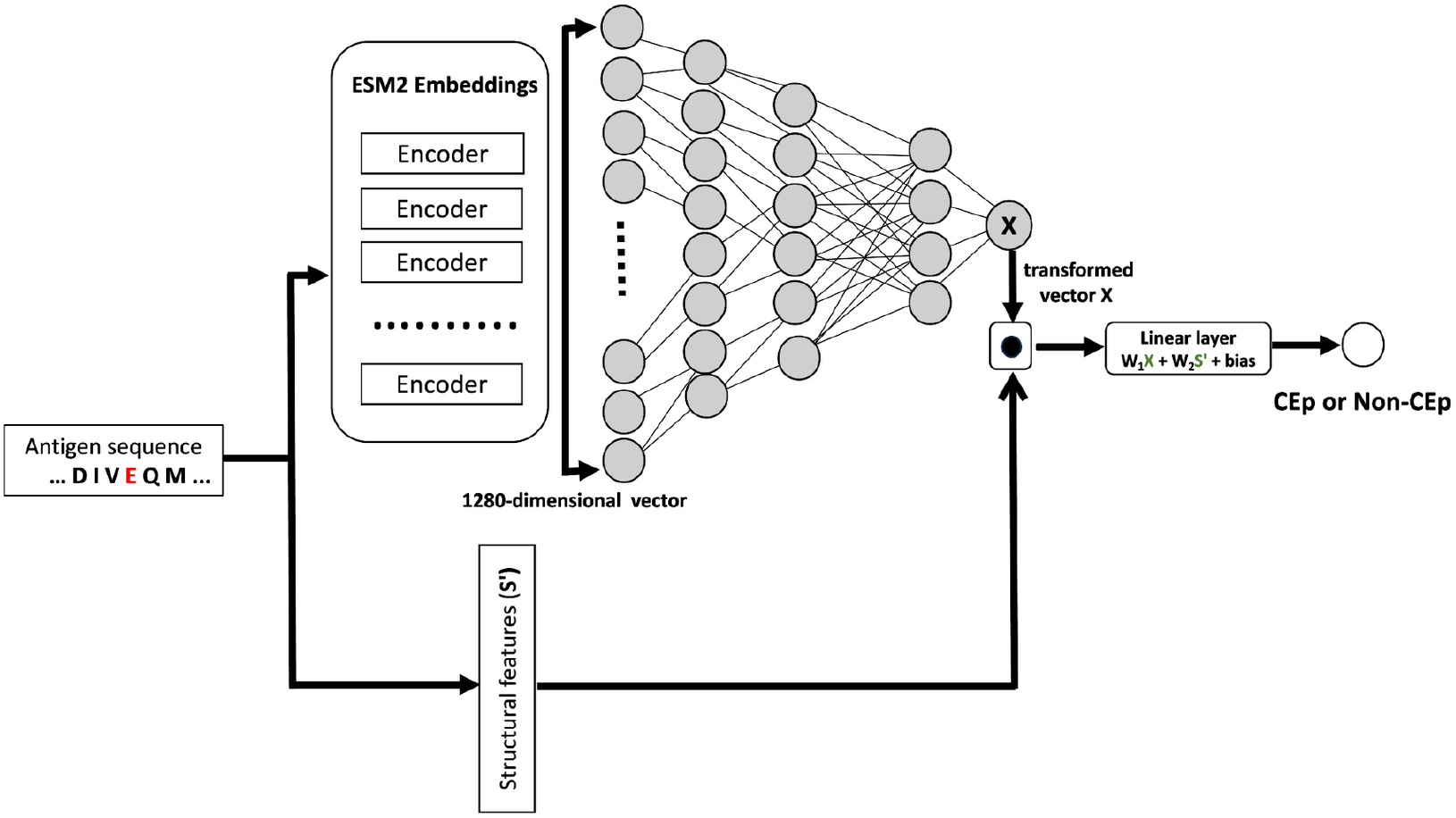
Model architecture II uses a deep concatenation of structural features and transformed ESM-2 embeddings in the final layer of the neural network to produce a probability score that classifies each residue as either epitope or non-epitope.

To learn the weights of the model in both the architectures we used a 5-fold cross validation technique with an outer testing loop and an inner validation loop for training. In the outer loop, to obtain reliable and stable predictions and to avoid over-fitting and biases in the training set, we split 901 complexes, into five sets randomly. Four sets out of five were used for training. This training set was split in the ratio 85:15 and used for training model and hyper parameter optimization. This step was performed 5 times so that we have 5 trained models and 5 test sets. The metrics from these five test sets corresponding to their models were averaged to get the final metric. We performed extensive hyperparameter tuning on the different number of layers, number of nodes in each layer, type of optimizer, learning rate to ensure a good, learned model (Table S2). Regularization techniques such as dropout layers, early stopping was also implemented to mitigate the over-fitting issues during the learning process of the model.

## 3. Results

### 3.1 Conformational epitope prediction

We trained various models on conformational epitope data using two architectures and various feature combinations. We evaluated our models on the independent test set consisting of 45 antigens. For the final prediction for each datapoint, we calculated the mean of the predicted scores from the 5 models which were trained using the nested 5-fold cross validation technique. The results in Figure 3 show that architecture II, that used feature transformation and concatenation technique, performed better than architecture I that used naive concatenation with the same set of features. Architecture II that uses ESM-2 and CN as features, performs the best with area under the receiver operating characteristic curve (AUROC) of 0.813 and area under precision-recall curve (AUPRC) of 0.445. Our best model performs better than Discotope3.0 (AUROC 0.755 and AUPRC 0.180) which is a state-of-the-art epitope prediction tool. Different conformational epitope prediction models using different feature combinations and architectures developed in this study and their metrics are shown in Table S1.

**Figure 3.**
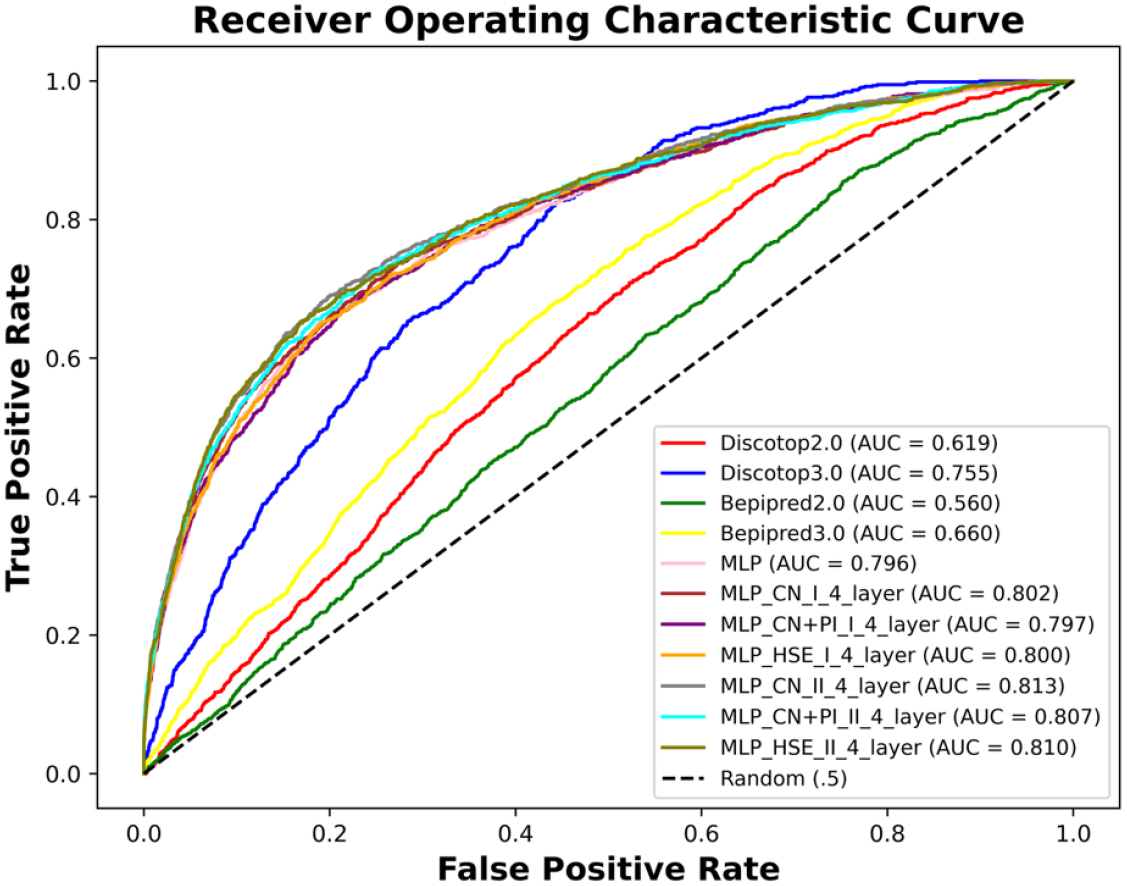
Comparison of models based on ROC-curves

We also conducted experiments by replacing ESM2 embeddings with structure aware SaProt (Su et al., 2024) embeddings to check if the later performs better. The results in Table S3 show that the AUROC of the model using only SaProt embeddings is 0.655 while AUROC of model using only ESM2 embeddings is 0.798. A model that uses SaProt embeddings combined with structural features performs better than the model that uses only SaProt embeddings, underscoring the importance of structural features used in this study for predicting the epitopes. In these experiments too, architecture II demonstrates higher performance compared to architecture I, indicating that exploring different feature combination methods beyond feature concatenation may be beneficial.

### 3.2 Linear epitope prediction

Most of the conformational epitope prediction tools use structural features, however linear epitope predictors have so far mostly used sequence-based features except Ellipro (PI) (Ponomarenko et al., 2008). It has been shown that structural features can be important for predicting linear epitopes as well (Barlow et al., 1986). Thus, we trained a linear epitope predictor using the same set of combination features (ESM-2, PI, CN, HSE) as we did for the conformational epitopes. The linear epitope train, validation and test data was prepared as detailed in the data section. Similar architectures and feature combination methods were used as for conformational epitope models. As shown in Figure 4, our best linear model that uses deep transformation and concatenation of ESM-2 features with structural features CN and PI, has AUROC score of 0.78 and outperforms BepiPred3.0 and Epidope linear epitope predictors that have AUROC of 0.49 and 0.56 respectively. We also tested the performance of our best conformational epitope models trained on the conformational data on the linear epitope test data. As seen in Figure 4, models trained on linear epitope data perform better on the linear test dataset than the ones trained on the conformational data. Table S3 shows various metrics of linear epitope prediction models.

**Figure 4.**
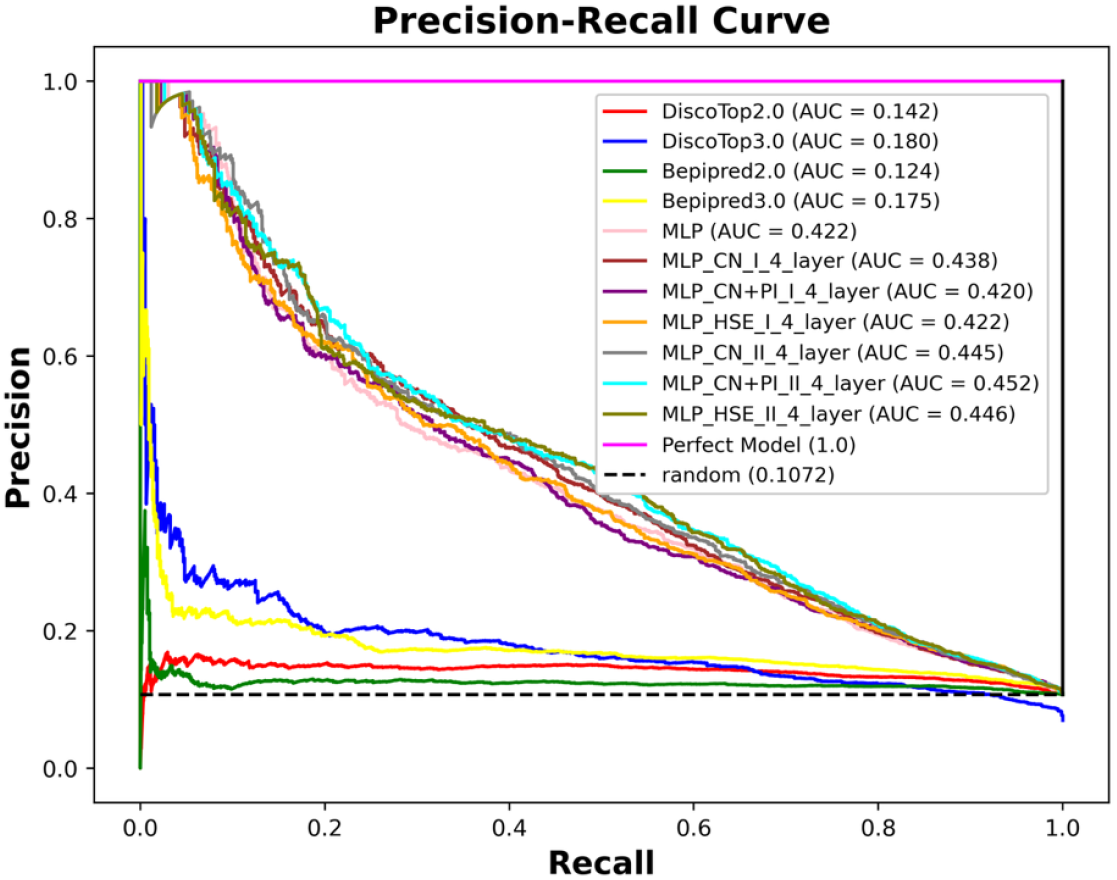
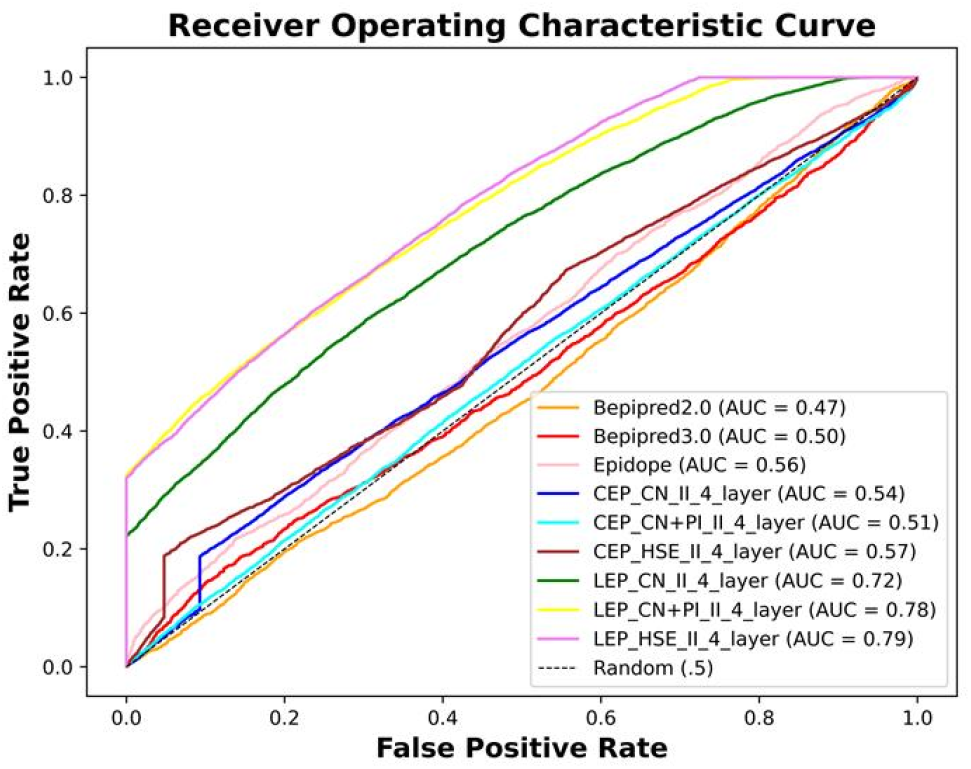
Comparison of models (trained and tested on linear epitope data) based on ROC-curves. Models with prefix CEP are conformational epitope prediction models tested on linear epitope test data and prefix LEP are Linear epitope prediction models trained and tested on linear epitope data.

### 3.3 ESM embeddings capture T-B reciprocity

Research indicates that T cells assist B cells in producing antibodies (Lanzavecchia, 1985). The binding of antibodies affects antigen processing, influenced by the spatial relationship between T-cell and B-cell epitopes. According to the immunodominance theory, these positions impact how dominant an immune response becomes (Biavasco and De Giovanni, 2022). CD4 T helper cells help select high-affinity B cell receptors within specific epitope environments, a process termed T-B reciprocity (Zhu et al., 2022). Essentially, B-cell epitopes near T-cell epitopes show increased immunogenicity. Based on these studies, Zhu et al. developed a model for linear epitope prediction that incorporated T-cell proximity as a feature to improve the prediction of immunogenic epitopes.

We know from previous studies that ESM-2 captures physicochemical properties of proteins, stores information on evolutionary contacts (Lin et al., 2022) and conservation (Marquet et al., 2022). Thus, we did attention analysis to check if T-B reciprocity was captured by vanilla ESM-2 weights. We reasoned that the pairwise residue importance for T and B epitope residues captured from the attention matrix should be statistically significant. For our analysis, we shortlisted 156 antigens from IEDB which had both T-cell and B-cell epitope annotations. Thus, this analysis was restricted to known (true) epitopes only. We extracted the attention matrices for these antigens from the last layer of ESM-2 by averaging out the attention matrices across the 20-attention heads. Each element of the attention matrix of an antigen represents the degree of importance or focus a residue of the antigen should give to other residues. From the antigen attention matrix, we got attention scores for all known T epitopes. The T-cell epitope residue positions’ attention score vectors were sorted in descending manner and from these sorted vectors we obtained the top residues whose attention scores sum up to 0.5 as these positions were considered highly attended. It was observed that in 116 antigens at least one B-cell epitope was attended and in 18 antigens all the known B-cell epitopes were highly attended by the T-cell epitopes. There were 40 cases where none of the B-cell epitopes were attended by any T-cell epitope. To examine the difference between these two extremities, we predicted the scores for the B-cell epitopes of these 58 sequences from our model and ran a statistical t-test (“ttest_ind — SciPy v1.14.1 Manual,” n.d.) (Virtanen et al. 2020) on the predicted scores for these two sets. The calculated p-value for this test was seen to be 0.03, which shows that there is a significant difference between the representation of the scores between these two data samples as seen in Figure 5.

**Figure 5.**
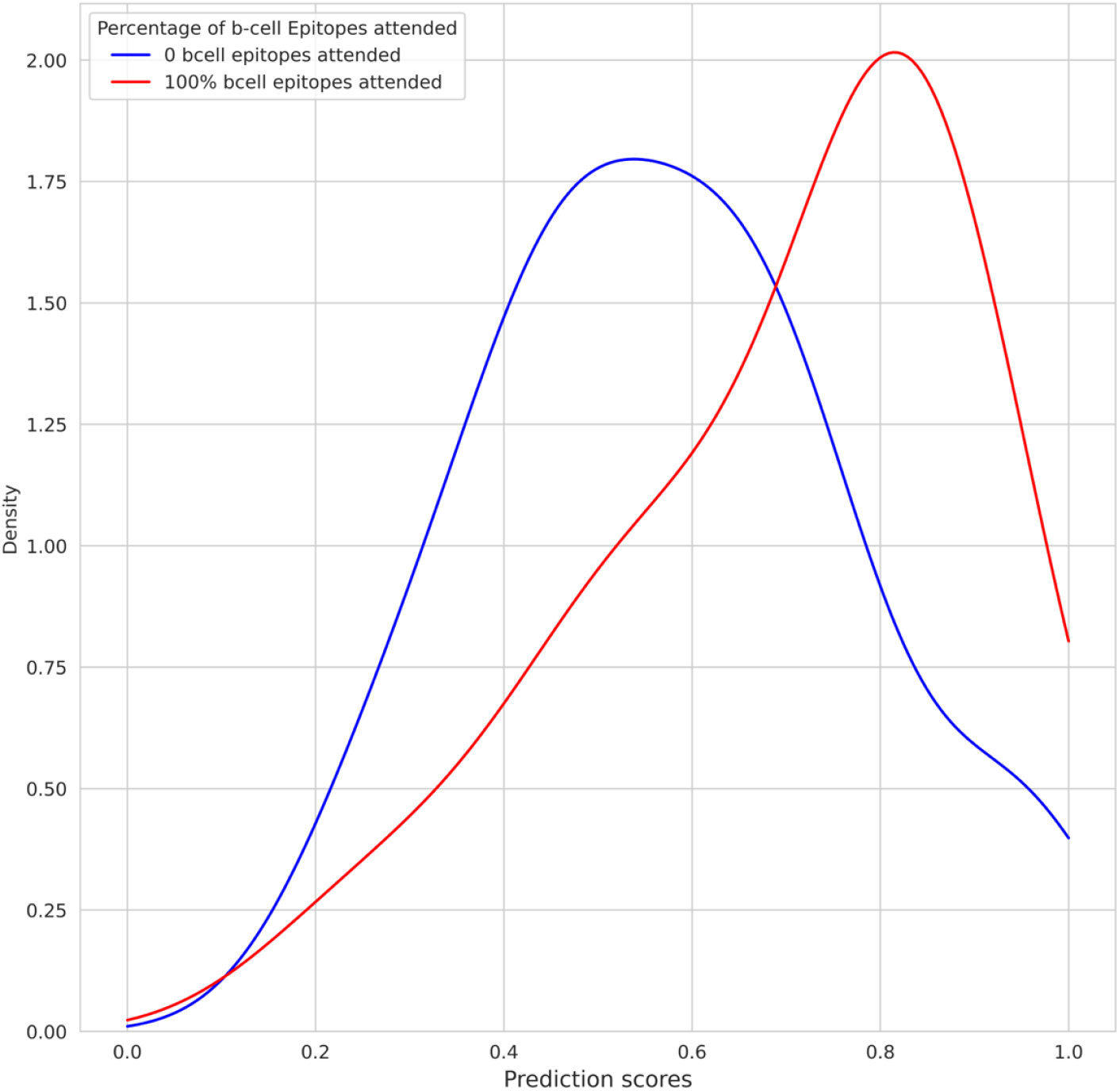
The distributions of the two extreme cases for the conformational epitope data have been shown here. The area under the red curve represents the distribution of the predicted scores for the sequences where all the B-cell epitopes were attended to by the T-cell epitopes. The area under the blue curve represents the distribution of the predicted scores for the sequences where none of the B-cell epitopes were attended to by the T-cell epitopes.

From these observations we can infer that the protein large language models likely capture some form of T-cell and B-cell epitope reciprocity, as seen with the help of attention. This finding is crucial as it allows the researcher to filter high scoring immunogenic (owing to T-B reciprocity) sequences for antigen design.

A similar attention based analysis was also done for linear B-cell epitopes. For this we took 254 antigen sequences for which both the B-cell and T-cell epitope information was known. It was observed that for certain sequences all the known B-cell epitopes were highly attended by the T-cell epitopes. Such cases were seen to be 31 out of the 254 antigens. There were 35 cases where none of the B-cell epitopes were attended by any T-cell epitope. To examine the difference between these two extremities, we predicted the scores for the B-cell epitopes of these 66 sequences and ran a statistical t-test on the predicted scores for these two sets. The calculated p-value for this test was seen to be 0.007, which shows that there is a significant difference between the scores between these two data samples (Figure 6).

**Figure 6.**
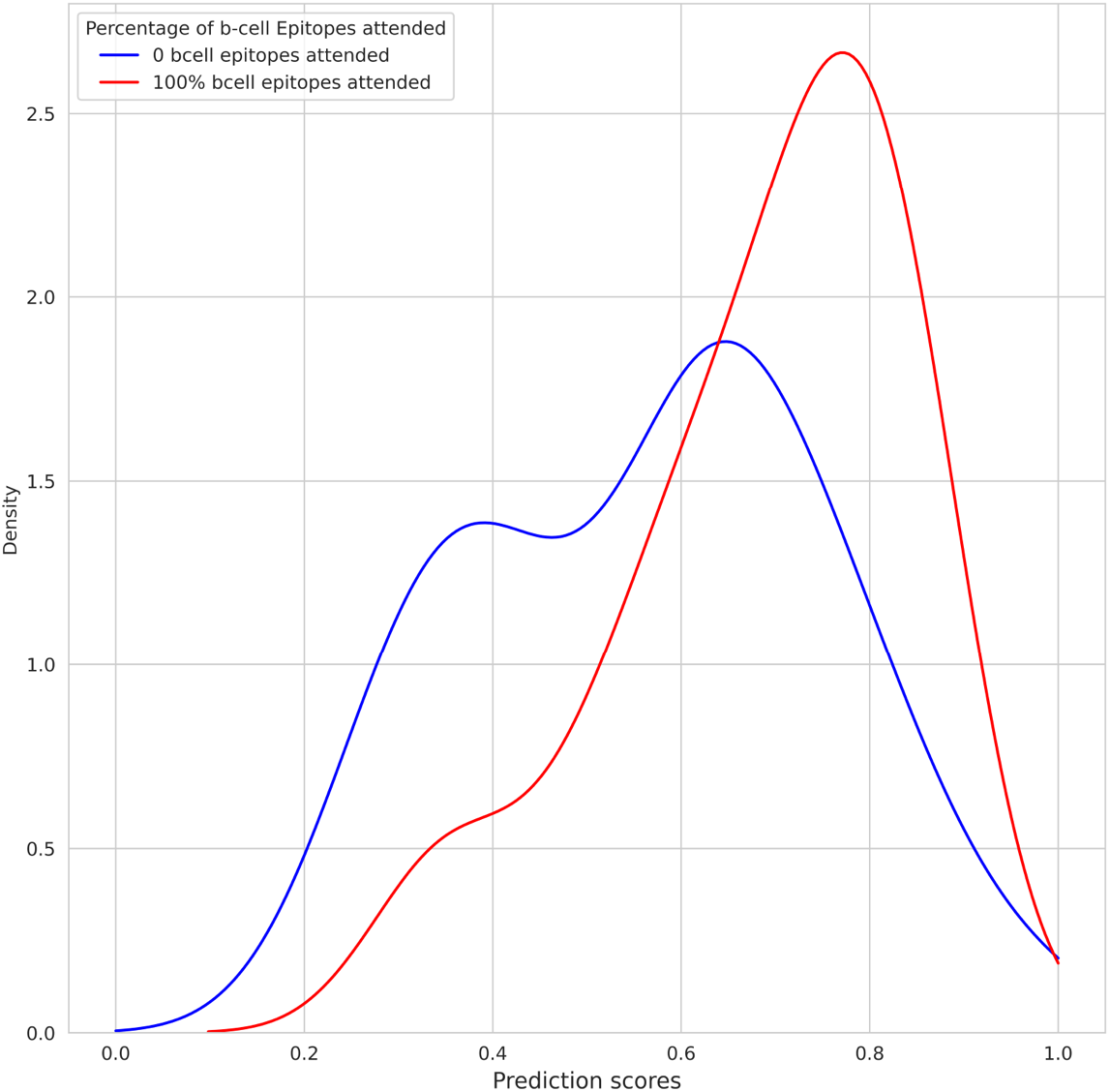
The distributions of the two extreme cases for the linear epitope data are shown here. The area under the red curve represents the distribution of the predicted scores for the sequences where all the B-cell epitopes were attended to by the T-cell epitopes. The area under the blue curve represents the distribution of the predicted scores for the sequences where none of the B-cell epitopes were attended to by the T-cell epitopes.

## 4 Discussion

In this study we demonstrate that epitope predictions from the ESM-2 model can be improved by combining structural features with the ESM-2 embeddings. We have also shown that the feature combination techniques can have an impact on the prediction accuracy of the models. We use two feature combination techniques: naive concatenation and feature transformation followed by concatenation. In the naive concatenation approach (Architecture I) features from ESM-2 embeddings (1280-dimension vector), are concatenated with structural features. This way of combining different feature sets is commonly seen to work well in most settings. This kind of approach ignores the fact that embeddings and structural features are from different spaces. Another approach of feature combination involved converting ESM-2 embeddings (1280 features) to a single transformed feature via layers of neural network. This transformed feature along with structural features is again passed through a linear layer to get a final weighted representation (Architecture II). We see from our analysis that Architecture II gives improved predictions as compared to Architecture I. We have used these two architectures for predicting both conformational and linear epitopes. We see improvement in both conformational and linear epitope prediction by combining ESM-2 embeddings with structural features. This is significant especially for linear epitopes as most algorithms use only sequence based features for linear epitope prediction except Ellipro (Ponomarenko et al., 2008) that uses protrusion index. Our best models (both linear and conformational epitopes) compare favorably with other state-of-the-art methods. We see that models with ESM-2 embeddings along with Contact number, Half Sphere Exposure and Protrusion Index as structural features are best, underscoring the importance of solvent exposure and protein shape for predicting epitopes. Our study also shows that T-B reciprocity is likely captured by ESM-2 and can be potentially used to differentiate high scoring immunogenic B epitopes that are highly attended by T epitopes from high scoring antigenic B epitopes.

## Author Contributions

S.R. Conceptualization and study design; R. Sajeed. and S.P. Implementation; R. Sajeed and S.P. Data processing; R. Sajeed Carried out all experiments; S.R. R. Sajeed, S.P and R. Srinivasan Analysis; S.R and R. Sajeed Wrote the original draft; SR and R. Srinivasan Supervision; SR, R. Srinivasan reviewed the manuscript

## Acknowledgements

Authors thank their colleague Dinesh Joshi for helpful discussions

## References

Bahai, A., Asgari, E., Mofrad, M.R.K., Kloetgen, A., McHardy, A.C., 2021. EpitopeVec: linear epitope prediction using deep protein sequence embeddings. Bioinformatics 37, 4517–4525. 10.1093/bioinformatics/btab467

Barlow, D.J., Edwards, M.S., Thornton, J.M., 1986. Continuous and discontinuous protein antigenic determinants. Nature 322, 747–748. 10.1038/322747a0

Biavasco, R., De Giovanni, M., 2022. The Relative Positioning of B and T Cell Epitopes Drives Immunodominance. Vaccines 10, 1227. 10.3390/vaccines10081227

Bio.PDB package — Biopython 1.75 documentation [WWW Document], n.d. URL https://biopython.org/docs/1.75/api/Bio.PDB.html (accessed 11.21.24).

Burley, S.K., Berman, H.M., Kleywegt, G.J., Markley, J.L., Nakamura, H., Velankar, S., 2017. Protein Data Bank (PDB): The Single Global Macromolecular Structure Archive. Methods Mol Biol 1607, 627–641. 10.1007/978-1-4939-7000-1_26

Clifford, J.N., Høie, M.H., Deleuran, S., Peters, B., Nielsen, M., Marcatili, P., 2022. BepiPred‐3.0: Improved B‐cell epitope prediction using protein language models. Protein Science : A Publication of the Protein Society 31, e4497. 10.1002/pro.4497

da Silva, B.M., Myung, Y., Ascher, D.B., Pires, D.E.V., 2022. epitope3D: a machine learning method for conformational B-cell epitope prediction. Brief Bioinform 23, bbab423. 10.1093/bib/bbab423

Desta, I.T., Kotelnikov, S., Jones, G., Ghani, U., Abyzov, M., Kholodov, Y., Standley, D.M., Beglov, D., Vajda, S., Kozakov, D., 2023. The ClusPro AbEMap web server for the prediction of antibody epitopes. Nature protocols 18, 1814. 10.1038/s41596-023-00826-7

Høie, M.H., Gade, F.S., Johansen, J.M., Würtzen, C., Winther, O., Nielsen, M., Marcatili, P., 2024. DiscoTope-3.0: improved B-cell epitope prediction using inverse folding latent representations. Frontiers in Immunology 15, 1322712. 10.3389/fimmu.2024.1322712

Hou, Q., Stringer, B., Waury, K., Capel, H., Haydarlou, R., Xue, F., Abeln, S., Heringa, J., Feenstra, K.A., 2021. SeRenDIP-CE: sequence-based interface prediction for conformational epitopes. Bioinformatics 37, 3421–3427. 10.1093/bioinformatics/btab321

Ivanisenko, N.V., Shashkova, T.I., Shevtsov, A., Sindeeva, M., Umerenkov, D., Kardymon, O., 2024. SEMA 2.0: web-platform for B-cell conformational epitopes prediction using artificial intelligence. Nucleic Acids Research 52, W533–W539. 10.1093/nar/gkae386

Jespersen, M.C., Peters, B., Nielsen, M., Marcatili, P., 2017. BepiPred-2.0: improving sequence-based B-cell epitope prediction using conformational epitopes. Nucleic Acids Research 45, W24. 10.1093/nar/gkx346

Jumper, J., Evans, R., Pritzel, A., Green, T., Figurnov, M., Ronneberger, O., Tunyasuvunakool, K., Bates, R., Žídek, A., Potapenko, A., Bridgland, A., Meyer, C., Kohl, S.A.A., Ballard, A.J., Cowie, A., Romera-Paredes, B., Nikolov, S., Jain, R., Adler, J., Back, T., Petersen, S., Reiman, D., Clancy, E., Zielinski, M., Steinegger, M., Pacholska, M., Berghammer, T., Bodenstein, S., Silver, D., Vinyals, O., Senior, A.W., Kavukcuoglu, K., Kohli, P., Hassabis, D., 2021. Highly accurate protein structure prediction with AlphaFold. Nature 596, 583–589. 10.1038/s41586-021-03819-2

Krawczyk, K., Liu, X., Baker, T., Shi, J., Deane, C.M., 2014. Improving B-cell epitope prediction and its application to global antibody-antigen docking. Bioinformatics 30, 2288. 10.1093/bioinformatics/btu190

Kringelum, J.V., Lundegaard, C., Lund, O., Nielsen, M., 2012. Reliable B Cell Epitope Predictions: Impacts of Method Development and Improved Benchmarking. PLoS Computational Biology 8, e1002829. 10.1371/journal.pcbi.1002829

Marquet C, Heinzinger M, Olenyi T, Dallago C, Erckert K, Bernhofer M, Nechaev D, Rost B. Embeddings from protein language models predict conservation and variant effects. Hum Genet. 2022 Oct;141(10):1629–1647. doi: 10.1007/s00439-021-02411-y. Epub 2021 Dec 30. PMID: 34967936; PMCID: PMC8716573.

Ponomarenko, J., Bui, H.-H., Li, W., Fusseder, N., Bourne, P.E., Sette, A., Peters, B., 2008. ElliPro: a new structure-based tool for the prediction of antibody epitopes. BMC Bioinformatics 9, 514. 10.1186/1471-2105-9-514

Ponomarenko, J., Papangelopoulos, N., Zajonc, D.M., Peters, B., Sette, A., Bourne, P.E., 2011. IEDB-3D: structural data within the immune epitope database. Nucleic Acids Res 39, D1164–D1170. 10.1093/nar/gkq888

Ren, J., Liu, Q., Ellis, J., Li, J., 2014. Tertiary structure-based prediction of conformational B-cell epitopes through B factors. Bioinformatics 30, i264–273. 10.1093/bioinformatics/btu281

Sanchez-Trincado, J.L., Gomez-Perosanz, M., Reche, P.A., 2017. Fundamentals and Methods for T- and B-Cell Epitope Prediction. Journal of Immunology Research 2017, 2680160. 10.1155/2017/2680160

Steinegger, M., Söding, J., 2017. MMseqs2: sensitive protein sequence searching for the analysis of massive data sets. 10.1101/079681

Su, Jin, Chenchen Han, Yuyang Zhou, Junjie Shan, Xibin Zhou, and Fajie Yuan. “SaProt: Protein Language Modeling with Structure-Aware Vocabulary.” bioRxiv, April 19, 2024. 10.1101/2023.10.01.560349.

Sweredoski, M.J., Baldi, P., 2008. PEPITO: improved discontinuous B-cell epitope prediction using multiple distance thresholds and half sphere exposure. Bioinformatics 24, 1459–1460. 10.1093/bioinformatics/btn199

Xia, J.-F., Zhao, X.-M., Song, J., Huang, D.-S., 2010. APIS: accurate prediction of hot spots in protein interfaces by combining protrusion index with solvent accessibility. BMC Bioinformatics 11, 174. 10.1186/1471-2105-11-174

Yuan, Z., 2005. Better prediction of protein contact number using a support vector regression analysis of amino acid sequence. BMC Bioinformatics 6, 248. 10.1186/1471-2105-6-248

Zhou, C., Chen, Z., Zhang, L., Yan, D., Mao, T., Tang, K., Qiu, T., Cao, Z., 2019. SEPPA 3.0—enhanced spatial epitope prediction enabling glycoprotein antigens. Nucleic Acids Research 47, W388. 10.1093/nar/gkz413

Zhu, J., Gouru, A., Wu, F., Berzofsky, J.A., Xie, Y., Wang, T., 2022. BepiTBR: T-B reciprocity enhances B cell epitope prediction. iScience 25. 10.1016/j.isci.2022.103764

Zeming Lin, Halil Akin, Roshan Rao, Brian Hie, Zhongkai Zhu, Wenting Lu, Nikita Smetanin, Robert Verkuil, Ori Kabeli, Yaniv Shmueli, Allan dos Santos Costa, Maryam Fazel-Zarandi, Tom Sercu, Salvatore Candido, Alexander Rives, 2022. Evolutionary-scale prediction of atomic level protein structure with a language model. Science 379,1123–1130 (2023). DOI:10.1126/science.ade2574ttest_ind — SciPy v1.14.1 Manual [WWW Document], n.d. URL https://docs.scipy.org/doc/scipy/reference/generated/scipy.stats.ttest_ind.html (accessed 11.21.24)

Lanzavecchia, A., 1985. Antigen-specific interaction between T and B cells. Nature 314, 537–539. 10.1038/314537a0

Virtanen P, Gommers R, Oliphant TE, et al (2020) SciPy 1.0: fundamental algorithms for scientific computing in Python. Nat Methods 17:261–272. 10.1038/s41592-019-0686-2

Cock, P.J.A. et al. Biopython: freely available Python tools for computational molecular biology and bioinformatics. Bioinformatics 2009 Jun 1; 25(11) 1422–3 10.1093/bioinformatics/btp163

