## Supplementary Information for "An improved deep learning model for immunogenic B epitope prediction"

TCS Research, Tata Consultancy Services

**Table S1:** Comparison of the models trained on new updated data with the 4Å test set on the above-mentioned metrics (CN: Contact number, CNw : Contact Number with a window length of 9 around the residue, PI: Protrusion index, HSE: Half-sphere exposure):

| Model | Accuracy | Precision | Recall | AUCROC | Balanced Accuracy | AUPRC |
|---|---|---|---|---|---|---|
| Bepipred2 | 0.573 | 0.124 | 0.492 | 0.56 | 0.537 | 0.124 |
| Discotope2 | 0.822 | 0.152 | 0.143 | 0.619 | 0.523 | 0.142 |
| Bepipred3 | 0.715 | 0.169 | 0.425 | 0.66 | 0.587 | 0.176 |
| DiscoTope3 | 0.810 | 0.154 | 0.383 | 0.755 | 0.612 | 0.180 |
| SEMA-1Dv2.0 | 0.596 | 0.133 | 0.505 | 0.594 | 0.556 | 0.156 |
| SEMA-3D* | 0.521 | 0.100 | 0.737 | 0.667 | 0.621 | 0.130 |
| Random Forest Classifier with structural features | 0.727 | 0.190 | 0.472 | 0.669 | 0.615 | 0.191 |
| MLP_S* | 0.722 | 0.24 | 0.723 | 0.796 | 0.722 | 0.438 |
| MLP | 0.701 | 0.225 | 0.736 | 0.798 | 0.716 | 0.424 |
| MLP_5-fold_4_layer_CN_I | 0.7 | 0.23 | 0.76 | 0.8 | 0.73 | 0.44 |
| MLP_5-fold_4_layer_PI_I | **0.76** | **0.27** | 0.71 | **0.81** | **0.74** | **0.44** |
| MLP_5-fold_4_layer_HSE_I | 0.69 | 0.22 | 0.76 | 0.8 | 0.72 | 0.42 |
| MLP_5-fold_4_layer_CN+PI_I | 0.69 | 0.22 | 0.75 | 0.8 | 0.72 | 0.42 |
| MLP_5-fold_4_layer_CN+HSE_I | 0.66 | 0.21 | 0.77 | 0.79 | 0.71 | 0.42 |
| MLP_5-fold_4_layer_PI+HSE_I | 0.71 | 0.23 | 0.73 | 0.8 | 0.72 | 0.42 |
| MLP_5-fold_4_layer_CN+PI+HSE_I | 0.67 | 0.21 | 0.76 | 0.79 | 0.71 | 0.41 |
| MLP_5-fold_4_layer_CN_II | 0.73 | **0.25** | 0.75 | **0.81** | **0.74** | **0.45** |
| MLP_5-fold_4_layer_PI_II | 0.72 | 0.24 | 0.74 | 0.8 | 0.73 | 0.44 |
| MLP_5-fold_4_layer_HSE_II | **0.75** | **0.26** | 0.72 | **0.81** | **0.74** | **0.45** |
| MLP_5-fold_4_layer_CN+PI_II | 0.72 | **0.24** | 0.75 | 0.81 | **0.73** | **0.45** |
| MLP_5-fold_4_layer_CN+HSE_II | 0.69 | 0.22 | 0.76 | 0.8 | 0.72 | 0.43 |
| MLP_5-fold_4_layer_PI+HSE_II | 0.74 | **0.25** | 0.72 | 0.81 | **0.73** | **0.44** |
| MLP_5-fold_4_layer_CN+PI+HSE_II | 0.72 | 0.23 | 0.72 | 0.79 | 0.72 | 0.38 |

Table S1 depicts the comparison of our models from Architecture I and Architecture II with the other state-of-the-art tools for epitope prediction. The suffix ‘I’ points towards to models of Architecture I and ‘II’ towards Architecture II. These comparisons have been done on our independent test set having 45 PDB complexes.

_S* represents the model which was trained on data without 5-fold cross validation as has been done for the others

* Here one pdb complex from our independent set couldn’t be predicted with the tool SEMA-3D. The metrics are shown for the remaining 44 complexes.

**Table S2**: Hyperparameters tried for the different models

| Model number | Number of Hidden layers (HL) | Hidden layer size | Dropout rate | Input Features |
|---|---|---|---|---|
| MLP | 3 | 512,250,10 | 0.6 | 1280 |
| MLP+(CN/PI) _I | 3 | 512,250,10 | 0.6 | 1281 |
| MLP+(HSE)_I | 3 | 512,250,10 | 0.6 | 1282 |
| MLP+(PI+CN) _I | 3 | 512,250,10 | 0.6 | 1282 |
| MLP+(CN+HSE/PI+HSE) _I | 3 | 512,250,10 | 0.6 | 1283 |
| MLP+(CN+PI+HSE) _I | 3 | 512,250,10 | 0.6 | 1284 |
| MLP+(CN/PI) _II | 3 | 512,250,10 | 0.6 | 1280 |
| MLP+(HSE)_II | 3 | 512,250,10 | 0.6 | 1280 |
| MLP+(PI+CN) _II | 3 | 512,250,10 | 0.6 | 1280 |
| MLP+(CN+HSE/PI+HSE) _II | 3 | 512,250,10 | 0.6 | 1280 |

| | | | | |
|---|---|---|---|---|
| MLP+(CN+PI+HSE) _II | 3 | 512,250,10 | 0.6 | 1280 |

Computational resources – CPU or GPU
Learning rate - 1e-3
Activation functions – ReLU (used between the hidden layers) and Sigmoid (after the final layer for the output)

**Table S3:** Models trained on LEP (LEP_MLP_single_fold_4_layer models)/CEP (CEP_MLP_5_fold_4_layer models) data and tested on LEP data

| **Model_name** | **Acc** | **Pre** | **Rec** | **AUROC** | **Bal_acc** | **TPR** | **TNR** | **FPR** | **FNR** | **AUPRC** | **f1-score** |
|---|---|---|---|---|---|---|---|---|---|---|---|
| LEP_MLP_single_fold_4_layer_CN_II | 0.69 | 0.90 | 0.62 | 0.74 | 0.73 | 0.62 | **0.85** | **0.15** | 0.37 | 0.86 | 0.74 |
| LEP_MLP_single-fold_4_layer_CN+PI_II | 0.78 | 0.91 | **0.78** | 0.75 | 0.77 | **0.78** | 0.76 | 0.24 | **0.21** | 0.88 | **0.84** |
| LEP_MLP_single-fold_4_layer_HSE_II | **0.74** | **0.90** | 0.71 | **0.77** | **0.76** | 0.71 | 0.81 | 0.18 | 0.29 | **0.87** | 0.80 |
| CEP_MLP_5-fold_4_layer_CN_II | 0.58 | 0.90 | **0.35** | 0.64 | 0.64 | **0.35** | 0.94 | 0.05 | **0.64** | 0.76 | **0.50** |
| CEP_MLP_5-fold_4_layer_CN+PI_II | 0.54 | **0.90** | 0.22 | **0.60** | **0.59** | 0.22 | **0.96** | **0.03** | 0.77 | 0.69 | 0.36 |
| CEP_MLP_5-fold_4_layer_HSE_II | **0.55** | 0.90 | 0.28 | 0.61 | 0.62 | 0.28 | 0.95 | 0.04 | 0.71 | 0.73 | 0.43 |
| BepiPred2.0 | 0.50 | 0.52 | 0.70 | 0.47 | 0.48 | 0.70 | 0.26 | 0.73 | 0.30 | 0.52 | 0.60 |
| BepiPred3.0 | 0.49 | 0.57 | 0.25 | 0.49 | 0.51 | 0.25 | 0.77 | 0.22 | 0.74 | 0.55 | 0.34 |
| Epidope | 0.55 | 0.56 | 0.78 | 0.56 | 0.53 | 0.78 | 0.27 | 0.72 | 0.21 | 0.60 | 0.65 |

Table S3 shows the different metrics for the models trained on linear epitope data and tested on linear epitope test data which have been prefixed as ‘LEP_MLP_’, and metrics of models trained on conformational epitope data and tested on linear epitope test data which have been prefixed as ‘CEP_MLP_’. We have compared these models with the prediction tools BepiPred2.0 and 3.0, and EpiDope tools.

**Table S4:** Models trained on embeddings from SaProt 650M parameter model for the conformational epitope data

| **Model** | **Accuracy** | **Precision** | **Recall** | **AUCROC** | **Balanced Accuracy** | **AUPRC** |
|---|---|---|---|---|---|---|
| SAProt_35M | 0.553 | 0.162 | 0.656 | 0.628 | 0.598 | 0.174 |

| | | | | | | |
|---|---|---|---|---|---|---|
| SAProt_650M_PDB | 0.646 | 0.182 | 0.564 | 0.655 | 0.610 | 0.181 |
| SAProt_650M_PDB(+CN)_I | 0.600 | 0.172 | 0.620 | 0.650 | 0.609 | 0.182 |
| SAProt_650M_PDB(+PI)_I | 0.590 | 0.172 | 0.643 | 0.651 | 0.612 | 0.184 |
| SAProt_650M_PDB(+CN+PI)_I | 0.530 | 0.160 | 0.692 | 0.635 | 0.599 | 0.176 |
| SAProt_650M_PDB(+CN+PI+HSE)_I | 0.448 | 0.152 | 0.796 | 0.642 | 0.598 | 0.181 |
| SAProt_650M_PDB(+CN)_II | 0.442 | 0.152 | 0.803 | 0.640 | 0.598 | 0.176 |
| SAProt_650M_PDB(+PI)_II | 0.653 | 0.176 | 0.520 | 0.648 | 0.595 | 0.178 |
| SAProt_650M_PDB(+HSE)_II | 0.688 | 0.190 | 0.496 | 0.672 | 0.605 | 0.214 |
| SAProt_650M_PDB(+CN+PI)_II | 0.482 | 0.160 | 0.792 | 0.654 | 0.616 | 0.185 |
| SAProt(+CN+PI+HSE)_II | 0.702 | 0.201 | 0.503 | 0.674 | 0.616 | 0.218 |

Table S4 shows the metrics for the models trained on the conformational epitope data taking the SaProt 650M parameter model embeddings and using the structural feature combinations as well for both the architectures

**Figure S1**: Frequency distribution of the attended B-cell epitopes by the T-cell epitopes for the 156 complexes. Here the x-axis shows what percentage of the known B-cell epitopes are attended and the y-axis shows how many PDB complexes show this trend. Eg. For the first bar which shows 40 units, it shows that 40 PDB

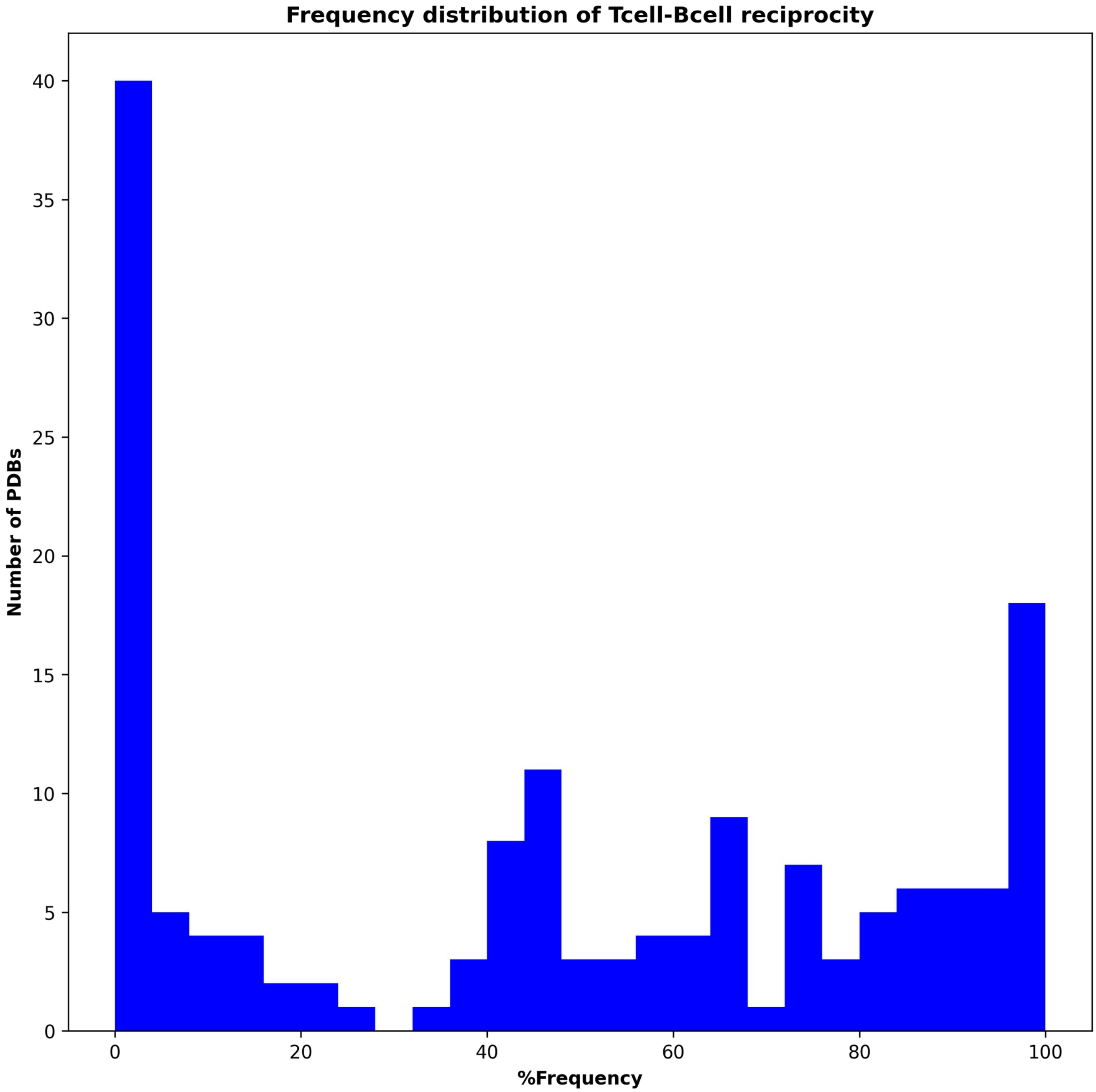


complexes have cases where the T-cell epitopes have attended to 0% of the known T-cell epitopes.
